# Correlated transcriptional responses provide insights into the synergy mechanisms of the furazolidone, vancomycin and sodium deoxycholate triple combination in *Escherichia coli*

**DOI:** 10.1101/2021.07.13.452286

**Authors:** Catrina Olivera, Murray Cox, Gareth J. Rowlands, Jasna Rakonjac

**Author notes:** Address correspondence to: Jasna Rakonjac. Department of Biology, University of Copenhagen, Copenhagen, Denmark (Catrina Olivera).

## Abstract

Effective therapeutic options are urgently needed to tackle antibiotic resistance. Furazolidone (FZ), vancomycin (VAN), and sodium deoxycholate (DOC) show promise as their combination can synergistically inhibit the growth of, and kill, multidrug-resistant Gram-negative bacteria that are classified as critical priority by the World Health Organization. Here, we investigated the mechanisms of action and synergy of this drug combination using a transcriptomics approach in the model bacterium *Escherichia coli*. We show that FZ and DOC elicit highly similar gene perturbations indicative of iron starvation, decreased respiration and metabolism, and translational stress. In contrast, VAN induced envelope stress responses, in agreement with its known role in peptidoglycan synthesis inhibition. FZ induced the SOS response consistent with its DNA damaging effects, but we demonstrate that using FZ in combination with the other two compounds enables use of lower dosages and largely mitigates its mutagenic effects. Based on the gene expression changes identified, we propose a synergy mechanism where the combined effects of FZ, VAN, and DOC amplify damage to Gram-negative bacteria while simultaneously suppressing antibiotic resistance mechanisms.

**Importance:** Synergistic combinations of existing antibacterials against multidrug-resistant “superbugs” are an alternative strategy to costly and arduous development of novel antibacterial molecules. The synergistic combination of nitrofurans, vancomycin and sodium deoxycholate shows promise in inhibiting and killing multidrug-resistant Gram-negative bacteria. We examined the mechanism of action and synergy of these three antibacterials, and proposed a mechanistic basis for their synergy. Our results highlight much needed mechanistic information necessary to advance this combination as a potential therapy.

## Introduction

Antimicrobial resistance is one of the biggest public health crises at present. With the traditional discovery and development of new antibiotics unable to keep pace with the emergence of resistance (1), alternative strategies are urgently needed to tackle multidrug-resistant bacteria. One promising approach is combining two or more drugs, especially if they are synergistic or have an enhanced combined effect (2, 3). Synergistic combinations can lead to better pathogen clearance, may slow down or prevent resistance development, and can lower the doses needed for each of the components, which in turn can mitigate adverse effects (3, 4). Repurposing existing drugs approved for human use can also be a faster way of bringing new therapies into the clinic in comparison to the development of novel antibacterial compounds (5).

Our recent studies have demonstrated the synergistic interaction of the existing antibiotics nitrofurans and vancomycin (VAN) with the secondary bile salt sodium deoxycholate (DOC) (6). In terms of efficacy and dose-reduction, we have shown that combining these three antibacterials is superior to the previously reported double combination synergy of nitrofuran and DOC (7) or nitrofuran and VAN (8). The triple combination is synergistic against a range of Gram-negative bacteria, including the critical priority pathogens, carbapenem-resistant *Enterobacteriaceae* and *Acinetobacter baumannii* (6, 9). We have characterized the nitrofuran, VAN, and DOC synergy *in vitro*, although the mechanism of synergy remains unknown.

Nitrofurans and DOC have variable effects on Gram-negative bacteria, but their exact mechanisms of action are not fully understood. Nitrofurans are prodrugs (10) whose reactive intermediates were reported to damage DNA, induce oxidative stress, and inhibit translation (11–14). On the other hand, the effects of DOC include DNA damage, oxidative stress, protein aggregation, and membrane damage (15–17). In contrast to FZ and DOC, VAN’s peptidoglycan synthesis inhibition in Gram-positive bacteria is well-characterized (18). However, since VAN is not used in the therapy against Gram-negative bacteria due to high minimum inhibitory concentrations (*e.g*. >100-fold higher MIC in *E. coli* than Gram-positive bacteria (6)), its effects on this group of organisms are currently unknown. VAN is a large hydrophilic glycopeptide antibiotic (MW = 1,449.3 Da) and cannot readily diffuse through the outer membrane porins which restrict molecule entry up to ~600 Da. Zhou et al. (2015) have proposed that small amounts of VAN can nevertheless cross the outer membrane and enter *E. coli*, making it possible for VAN to be synergistic with trimethoprim and the nitrofuran antibiotic nitrofurantoin. Increasing evidence also points to VAN having the same target and mechanism of action in Gram-negative bacteria. For example, low temperature can compromise the outer membrane and sensitize *E. coli* to VAN, and this sensitivity can be reversed by introducing an *Enterococcus* VAN resistance gene cluster that alters the target of the VAN compound (19). Furthermore, breaching the outer membrane in *E. coli* by expression of leaky mutant secretin channels lowers the MIC of VAN to as low as that for Gram-positive bacteria, and ruling out the possibility that target differences are the reason for the high MIC in Gram-negative bacteria (20).

Gram-negative bacteria are normally inherently resistant to VAN and DOC (21, 22), but the enhanced efficacy of the combination provides an opportunity to expand the use of these normally Gram-positive-only antibacterials to Gram-negative bacteria (6). Additionally, the possibility of dose reduction could mitigate nitrofuran’s reported mutagenicity (23, 24). The triple combination, therefore, shows considerable potential as a viable antibacterial treatment option. Understanding the mechanistic bases of the synergy will help advance this combination into the development pipeline and inform the rational design of superior combinations that include any of these antibacterials. This study examined the mechanisms of action and synergy of the nitrofuran furazolidone (<u>F</u>Z), <u>V</u>AN, and <u>D</u>OC (FVD combination) using a transcriptomics approach in the Gram-negative model bacterium *E. coli*. We show that the FVD combination mitigates nitrofuran mutagenicity, and by identifying perturbed pathways, we propose mechanisms for the action and synergistic interactions of the FVD combination.

## Results

### Extensive transcriptional responses to the FZ, DOC, and Van combination

We conducted RNA-Seq analysis to investigate the transcriptional profile of *E. coli* in response to FZ, VAN, and DOC alone or combination (FVD). To prevent the transcriptome profile from being overwhelmed by stochastic expression of cell death genes and other transcriptomic changes unrelated to the drug perturbations, we applied the treatments at subinhibitory concentrations and short exposure times (*i.e*. IC_50_ for 1 h, see Materials and Methods). Clustering samples into distinct groups by principal component analysis demonstrated high reproducibility across replicates and showed that all treatments, except for VAN, had distinctive effects on the transcriptome profile (Fig. S1A). Compared to the no-antibacterial control, we identified >1200 differentially expressed genes (DEGs) in each of the treatments (Dataset S1), except for VAN, which only resulted in 17 DEGs. FVD resulted in 95 upregulated and downregulated DEGs not found in the single antibacterials (Fig. S1C), and all FVD’s enriched Gene Ontology (GO) terms overlapped with those of FZ and DOC (Dataset S2).

To gain insights and compare the biological processes affected by the single antibacterials and FVD combination, we performed k-means clustering of all the DEGs (Fig. 1A) coupled with GO term enrichment analysis (Fig. 1B) (Dataset S3). The most significantly altered gene clusters in FZ-treated *E. coli* compared to the control were members of the SOS response (Fig.1A, Cluster 3) and respiration (Cluster 5), while those clusters that were most highly altered by DOC were involved in iron import, translation, and amino acid transport and synthesis (Fig. 1A, Clusters 1 and 4). Particularly striking was the major overlap of gene perturbations by FZ and DOC, and that the FVD combination resulted in the same pattern of gene clusters dysregulation, albeit sometimes less pronounced (*i.e*. smaller fold change relative to the control). FZ’s, DOC’s, and FVD’s upregulated genes were involved in iron import (Fig. 2A), ribosome assembly and translation (Fig. 2B), whereas downregulated genes appear in the respiratory/electron transport chain (ETC) (Fig. 2C) and central carbon metabolism (Fig. 2D).

**Fig. 1.**
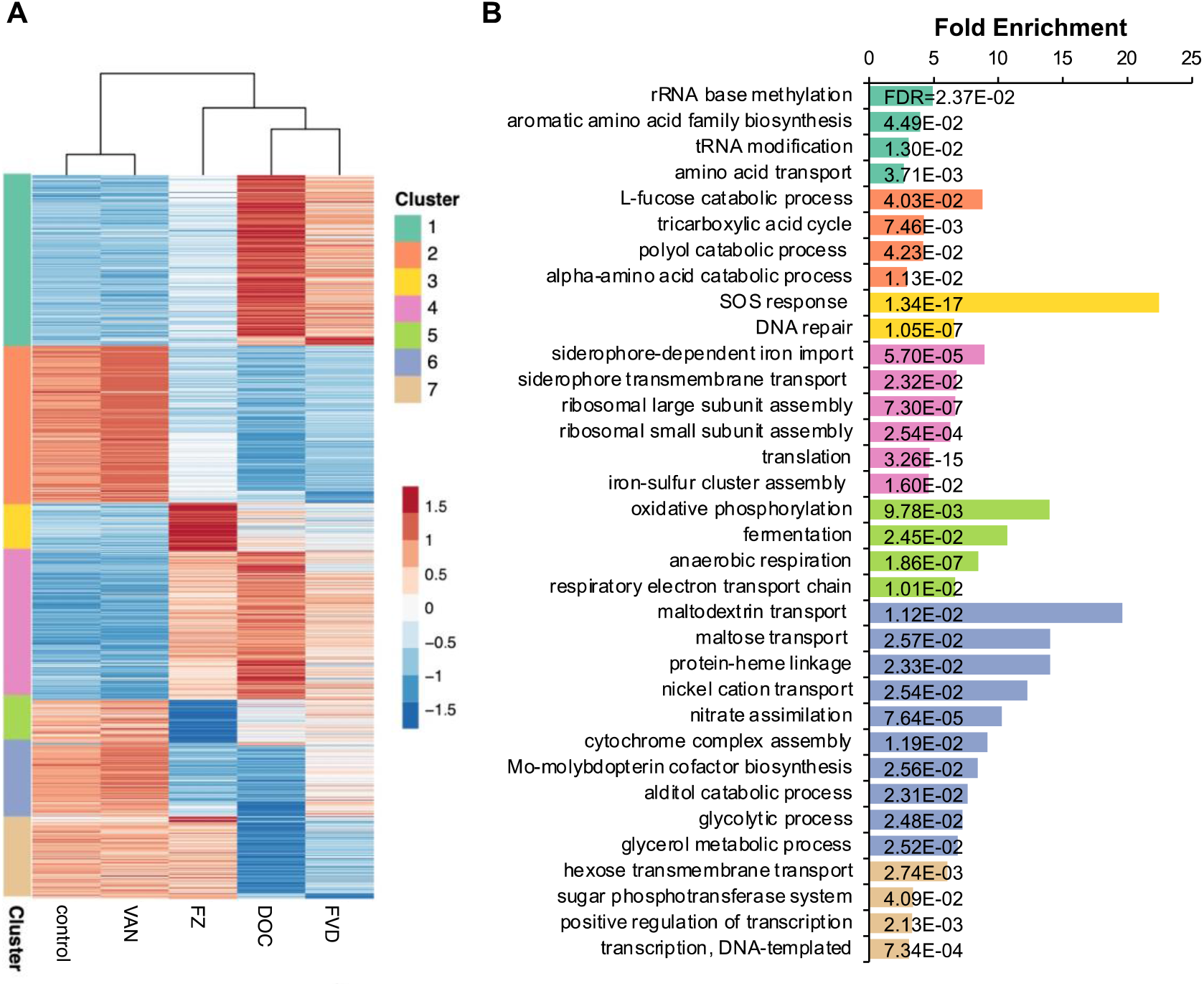
Heatmap of k-means clustered DEGs and GO term enrichment analysis. (A) Differentially expressed genes clustered into seven groups using k-means. Expression levels displayed were row-scaled regularized log-transformed normalized counts. (B) Each cluster was subjected to a biological process GO overrepresentation test, and the top enriched GO terms for each cluster are shown.

**Fig. 2.**
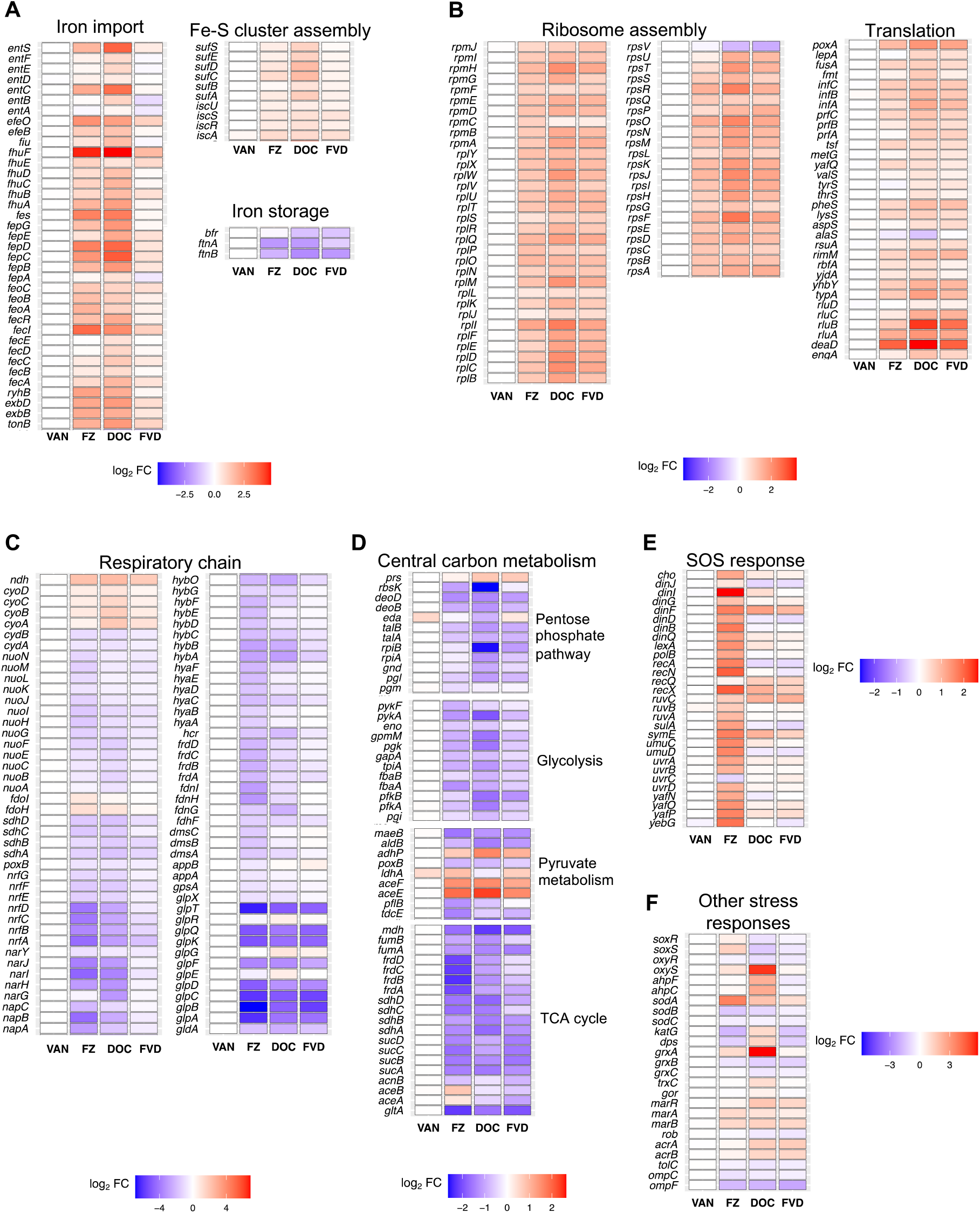
Differential expression of genes in *E. coli* K1508 treated with FZ, VAN, DOC, or the FVD combination. Expression (log_2_FC relative to no-antibacterial control) of genes involved in (A) iron homeostasis, (B) ribosome assembly and translation, (C) respiratory chain, (D) central carbon metabolism, (E) SOS response, and (F) oxidative and other antibiotic stress responses.

A closer look at the DEGs showed that the expression patterns in response to FZ and DOC indicate an iron starvation response; specifically, the upregulation of iron uptake, siderophore synthesis, Fe-S cluster assembly, and downregulation of iron storage and utilization (*e.g*. ETC, TCA cycle) (25, 26). The majority of these genes are either directly or indirectly controlled by the transcriptional repressor Ferric Uptake Regulator (Fur), which is inactivated by iron depletion (27–29). These results therefore suggest that FZ and DOC disturbs iron homeostasis either by signaling or triggering an iron starvation response. Other genes induced by FZ, DOC, or FVD include oxidative stress genes (*sodA*, *soxS*, *ahpF*, *ahpC*, *gor*, *grxA*) and those involved in increased antibiotic tolerance (Fig. 2F). The multiple antibiotic resistance genes *marA* and *marB*, of which the former is a master regulator of large number of genes involved in resistance, were both upregulated. Similarly, expression patterns for genes that encode efflux pumps and porins are suggestive of the inhibition of entry and accumulation of the antibiotics, such as the upregulation of efflux pump component genes *acrA* and *acrB* and downregulation of the outer membrane porin genes *ompC* and *ompF* (Fig. 2F).

### The FVD combination mitigates FZ-induced DNA damage

Genes involved in the SOS response were significantly upregulated in response to FZ, while other conditions, including the FVD combination, did not show the same level of SOS response gene upregulation (Fig. 2E), indicating the absence of severe DNA damage. To assess DNA damage levels induced by the antibacterials alone and in combination, we determined mutation frequencies in *E. coli* by quantifying mutant clones that gained resistance to rifampicin. Rifampicin is an RNA polymerase inhibitor, and resistance can arise through single base substitutions in the RNA polymerase gene *rpoB* (30). Under nonstress conditions (no-antibacterial control), the spontaneous mutation frequency of *E. coli* K1508 is around 7 mutants per 10^8^ cells (Fig. 3). Expectedly, given the DNA-damaging effects of FZ (11, 12), this number significantly increased upon FZ treatment (the mean frequency more than doubled). The other two single compounds (DOC and VAN) and the FVD combination, on the other hand, did not result in a significant change in rifampicin mutation frequency compared to the control. In comparison to the single antibacterials, the SOS gene induction (Fig. 2E) and rifampicin mutation frequency (Fig. 3) by FVD is considerably lower than those by FZ alone and more similar to DOC, suggesting that the use of the combination, in which the FZ concentration is lower than when used alone, can mitigate nitrofuran mutagenicity.

**Fig. 3.**
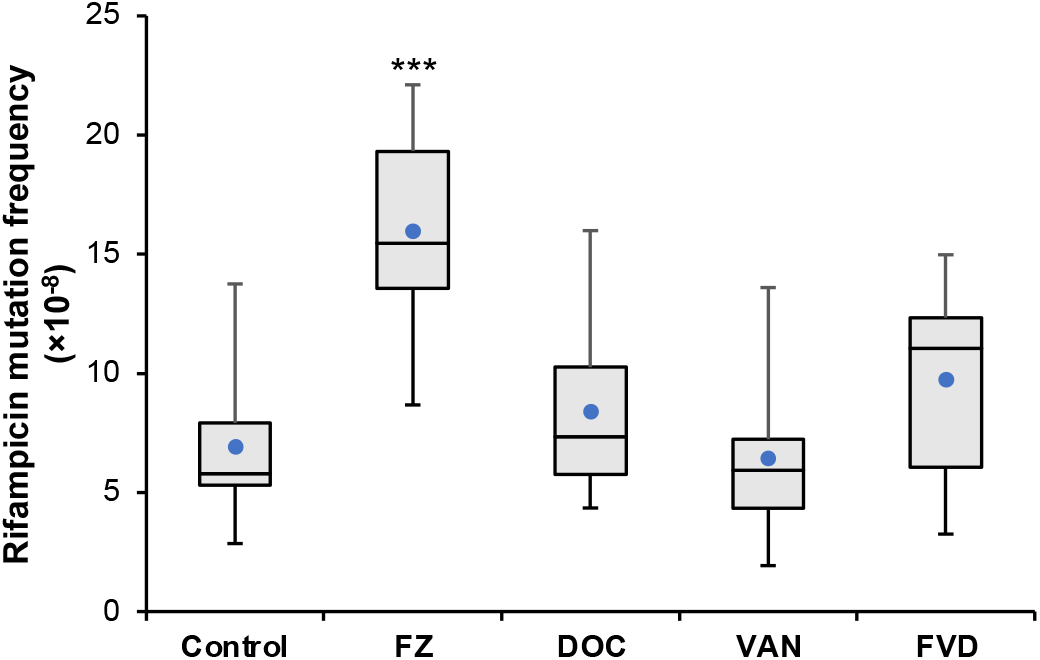
Rifampicin resistance mutation frequency in *E. coli* K1508 treated with FZ, DOC, VAN or the FVD combination. The mutation frequency of *E. coli* K1508 cells in the presence of antibacterials were determined by counting the colonies on rifampicin (100 μg/mL) plates after 24 h incubation. Box plots are derived from 9-11 biological replicates. Boxes show the interquartile range and the whiskers indicate the minimum and maximum values. The mean is indicated by a blue point. Statistical significance (Mann-Whitney U test of treatment vs control) is indicated by asterisks: ***, p < 0.001.

### FZ and DOC decelerate cellular respiration

Downregulation of genes encoding both aerobic and anaerobic ETC enzymes (Fig. 2C) prompted the assessment of physiological changes at the level of cellular respiration. Since oxygen is the major electron acceptor of the *E. coli* ETC (31), we investigated the overall effect on respiration by measuring oxygen consumption using a Clark-type oxygen electrode (Fig. 4). Exposure to IC_50_ FZ, DOC, and Van for 1 h significantly decreased the oxygen consumption rate, causing a 1.6-fold, 1.7-fold, and 1.15-fold decrease, respectively, compared to the no-antibacterial control. Similarly, the FVD combination reduced oxygen consumption by ~1.7-fold. The pattern of dysregulation of the aerobic ETC genes by FZ, DOC, and FVD potentially reflects an intracellular iron starvation signal (Fig. 2A), which is known to activate iron-sparing mechanisms (28). In this case, the non-iron utilising NADH dehydrogenase II (*ndh*) was possibly upregulated to compensate for the downregulation of iron-rich NADH dehydrogenase I (*nuoABCEFGHIJKLMN*) (Fig. 2C). In this assay, however, it was shown that the overall effect is a decrease in aerobic respiration.

**Fig. 4.**
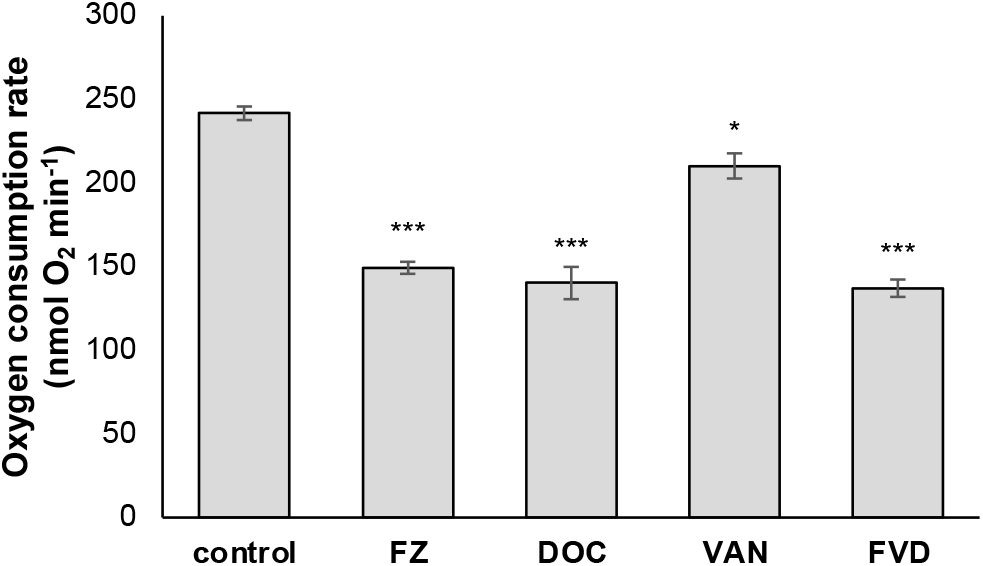
Effect of FZ, DOC, VAN, or the FVD combination on oxygen consumption. Oxygen consumption by *E. coli* K1508 was measured with a Clark-type closed-chamber oxygen electrode during exposure to IC_50_ FZ, DOC, and VAN alone and in combination. The cells were pre-treated with the drugs for 1 h, then oxygen consumption was measured for up to 5 min. Data show the mean of 3 replicates ± standard error of the mean. Statistical significance (Student’s t-test of treatment vs control) is indicated by asterisks as follows: *, *p* < 0.05; ***, *p* < 0.001.

### FZ and DOC dysregulate metal homeostasis

Metal homeostases in bacteria are highly interconnected (32). FZ and DOC dysregulation of genes involved in iron homeostasis prompted us to investigate the intracellular levels of iron and other essential metals (*e.g*. Mg, Cu, Ni, Zn, Mn) using inductively coupled plasma mass spectrometry (ICP-MS) analysis (Fig. 5). FZ treatment for 1 h resulted in a 1.8-fold decrease in iron and a 3-fold increase in manganese levels. On the other hand, DOC resulted in a reduction of magnesium, iron, and manganese and an increase in copper levels. Lastly, the effects of the FVD combination are the same as DOC, indicating that DOC drives most of the metal homeostasis changes. However, the fold changes caused by FVD are much more pronounced than those by DOC, reflecting the synergistic effect, such as the 18-fold decrease in magnesium (*vs*. 9-fold by DOC), 2-fold decrease in iron (*vs*. 1.6-fold by DOC), 5-fold decrease in manganese (*vs*. 2-fold by DOC) and a 6.5-fold increase in copper (*vs*. 2.5-fold by DOC) (Fig. 5). There was no significant change in the total intracellular zinc and nickel levels by any of the treatments, and VAN did not affect the total intracellular levels of any of the metals analysed in this study.

**Fig. 5.**
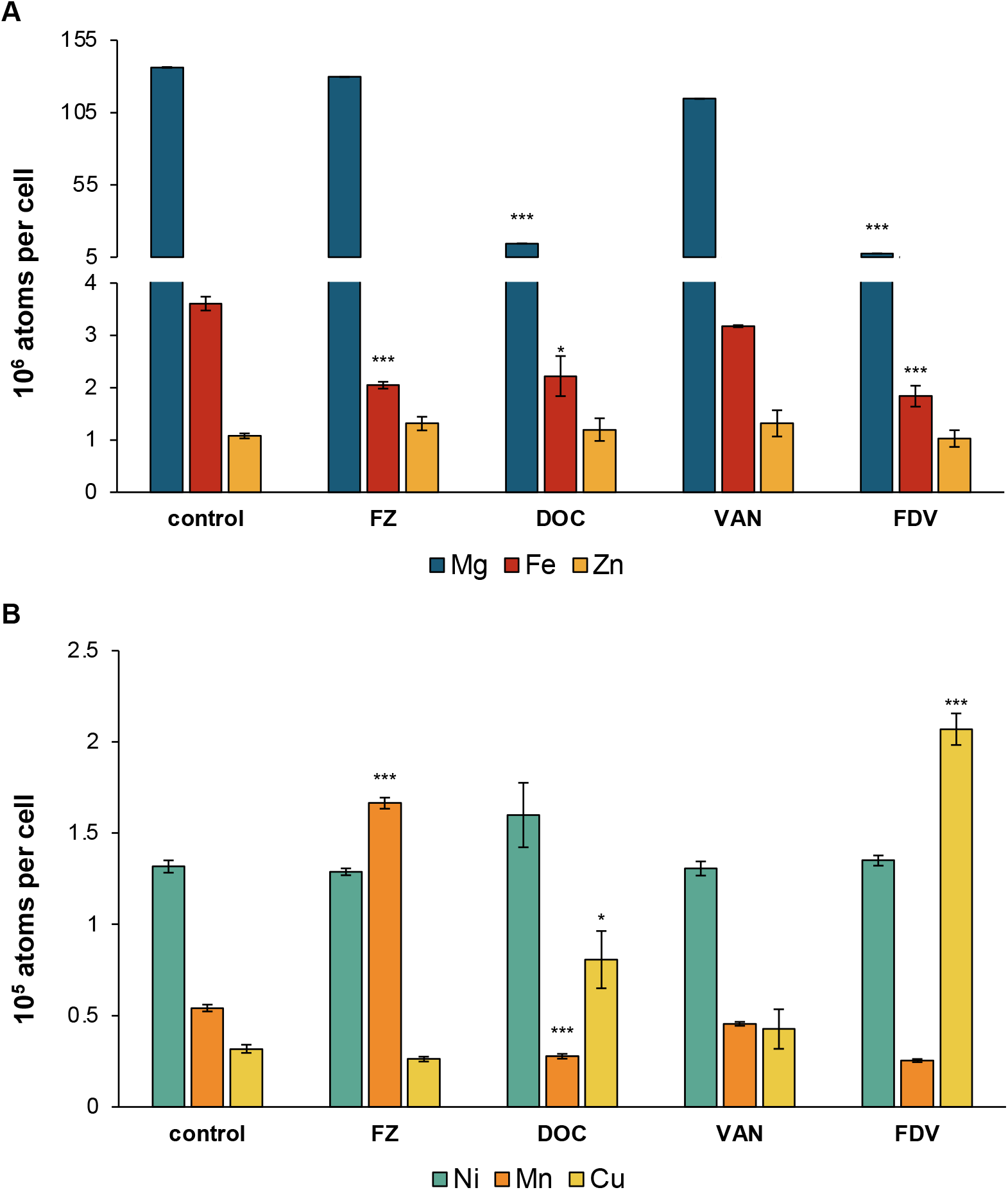
Intracellular metal levels after treatment with FZ, DOC, VAN or the FVD combination. Due to markedly different cellular concentrations, metals are shown in two graphs: (A) magnesium, iron and zinc (high levels), and (B) nickel, manganese and copper (low levels). Total intracellular metal concentrations were measured using ICP-MS in *E. coli* K1508 exposed to IC_50_ doses of FZ, DOC and VAN alone and the FVD combination for 1 h. Data show the mean of 3 replicates ± standard error of the mean. Statistical significance (Student’s t-test of treatment vs control) is indicated by asterisks: *, *p* < 0.05; ***, *p* < 0.001.

The observed changes in metal levels could partly explain the observed transcriptional response to FZ and DOC. For example, copper excess has been reported to degrade Fe-S clusters, block Fe-S cluster assembly, and stimulate an iron starvation response in *E. coli* and other bacteria (33–36). Therefore, copper toxicity could be a contributing factor that results in an iron starvation response to DOC or FVD. Similarly, during iron starvation or oxidative stress, manganese import is upregulated to replace iron as a cofactor in essential enzymes or prevent oxidative protein damage (37, 38), and could therefore explain increased manganese levels by FZ. Surprisingly, even though DOC induced the expression of oxidative stress genes of the OxyR regulon (*e.g*. *ahpC*, *ahpF*, *katG*, *dps*, *grxA*, *trxC*, *oxyS*) (Fig. 2F) indicating H_2_O_2_ stress (39), it caused a decrease in total intracellular manganese levels. Taken together, FZ and DOC, besides affecting iron homeostasis, also result in the dysregulation of other essential metals, including manganese, magnesium, and copper.

### Possible inhibition of SOS response by VAN

Most of the 17 DEGs triggered by Van were found to belong to multiple stress response regulons, most frequently those involved in envelope stress: Rcs, Cpx, and Bae (Fig. 6). Upregulation of these genes is consistent with the effects of peptidoglycan synthesis inhibitors that are effective against Gram*-*negative bacteria (40, 41) and supports a VAN mechanism of action identical to that in Gram-positive bacteria (42). Similarly, DOC and FVD upregulated most of the envelope stress genes induced by VAN (Fig. 6). This is not surprising since DOC is known to disrupt biological membranes (15, 17). It is possible that membrane disruption by DOC allows more Van to enter and exert its effect leading to synergy.

**Fig. 6.**
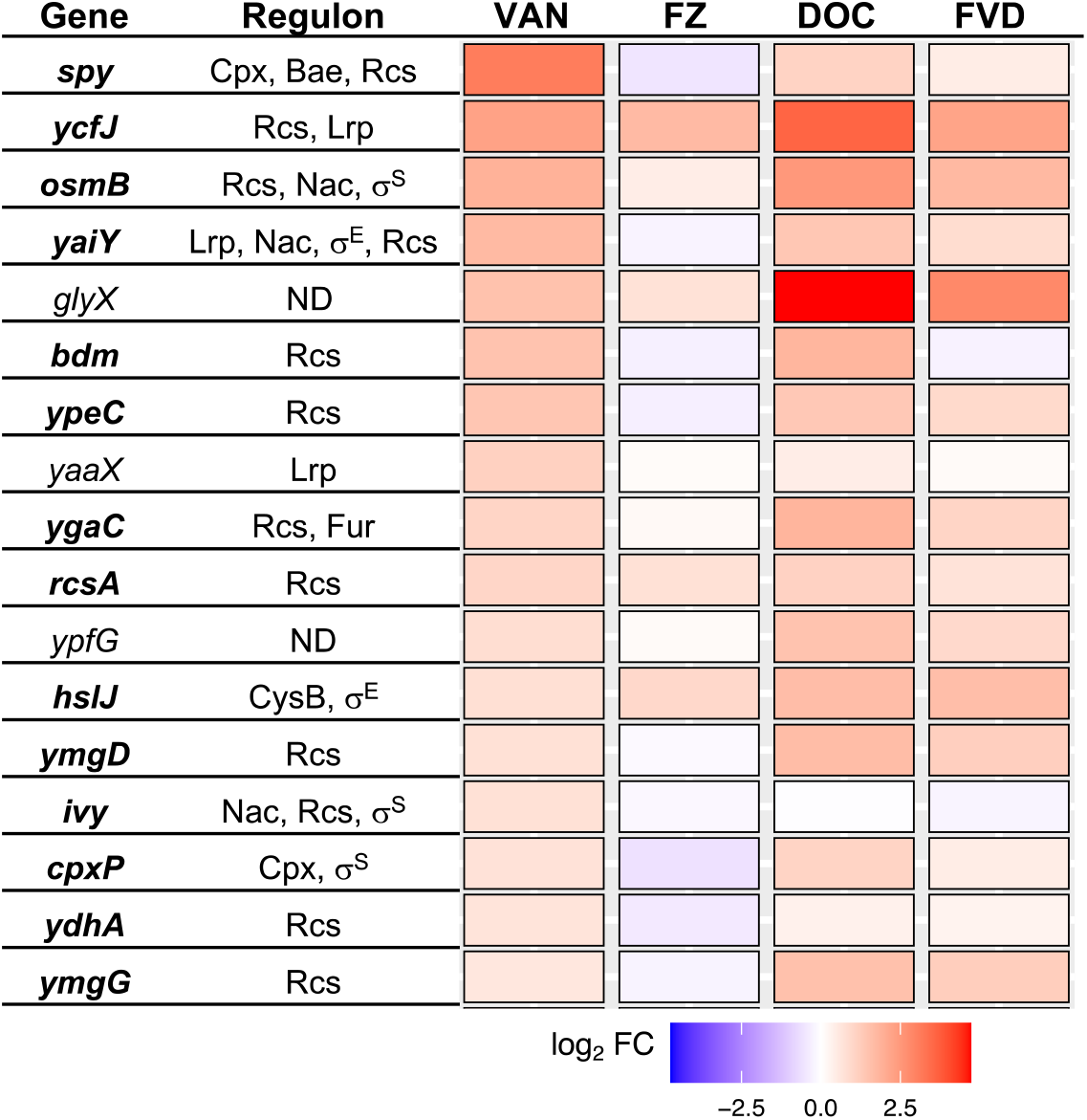
Differential expression of the 17 VAN DEGs across all treatments. Gene expression differences were presented as log_2_FC relative to no-antibacterial control. Regulators of each gene were derived from information in EcoCyc (https://ecocyc.org/), RegulonDB (http://regulondb.ccg.unam.mx/) and the literature (40, 43); genes with undetermined regulator are shown as ND. Genes involved in envelope stress are in **bold**.

To gain more insights into VAN effects on *E. coli*, we analysed the transcriptomics data by O’Rourke *et al*. (44), who investigated the transcriptional response of an *E. coli* strain with a compromised outer membrane to various antibiotics, including VAN (44). We performed biological process GO term enrichment of the significantly upregulated and downregulated genes by VAN (*i.e*. genes with FDR < 0.1 in Supplementary dataset S2 from ref (44)), but we did not use a fold change cut-off, as in that paper, so as not to omit any information from lowly expressed genes. Biological process GO term enrichment of significantly downregulated genes (FDR < 0.01, log_2_FC < 0) showed the SOS response to be overrepresented (Fig. S2). The SOS response genes downregulated by VAN in the O’Rourke *et al*. dataset are *umuC*, *umuD*, *uvrB*, *uvrC*, *uvrD*, *recN*, *dinB*, *dinG*, *polB*, *cho*, and *sulA*. Downregulation of these genes could thus explain the synergy between nitrofuran and VAN reported by Zhou *et al*. (8), who hypothesized that VAN must increase DNA-damaging effects when combined with DNA-damaging agents, such as nitrofurantoin or trimethoprim. If VAN exerts the same inhibition of SOS/DNA repair in wildtype *E. coli*, this effect will contribute to the triple synergy by amplifying the DNA-damaging effects of FZ and DOC, in such a way that it decreases DNA damage adaptation and survival through mutagenicity, which increases lethality.

If the SOS response plays a role in the synergy of FVD, for example through the inhibition of SOS response by VAN, deletion of *recA*, which makes *E. coli* unable to mount an SOS response (45), is expected to disrupt the FVD combination’s synergy mechanism, therefore decreasing the synergy in the mutant strain. Expectedly, *recA* deletion increased the susceptibility to FZ by 32-fold relative to the wildtype. Likewise, this deletion also decreased DOC MIC to 80000 μg/mL from more than 80000 μg/mL, while VAN MIC (250 μg/mL) remained unchanged. In the checkerboard assay to investigate the interaction of FZ, VAN, and DOC, deletion of *recA* caused a slight increase in the interaction index (FICI) of FVD (FICI < 0.22) compared to the wildtype (FICI < 0.13), indicating only a slight decrease in synergy (Fig. S3). For the two-drug interactions, only FZ and VAN combination showed a significant change in the FICI in the *recA* mutant (Fig. 7). The deletion of *recA* resulted in a shift to indifferent interaction (FICI = 1) instead of the synergy observed in the wildtype (FICI < 0.38) (Fig. 7A). Taken together, these findings support the hypothesis that the SOS response is an interacting point for the synergy between FZ and VAN. In terms of the triple combination synergy, however, the SOS response contributes to the synergy, but, given that deletion of *recA* still results in a synergistic interaction, other factors contributing to synergy are present.

**Fig. 7.**
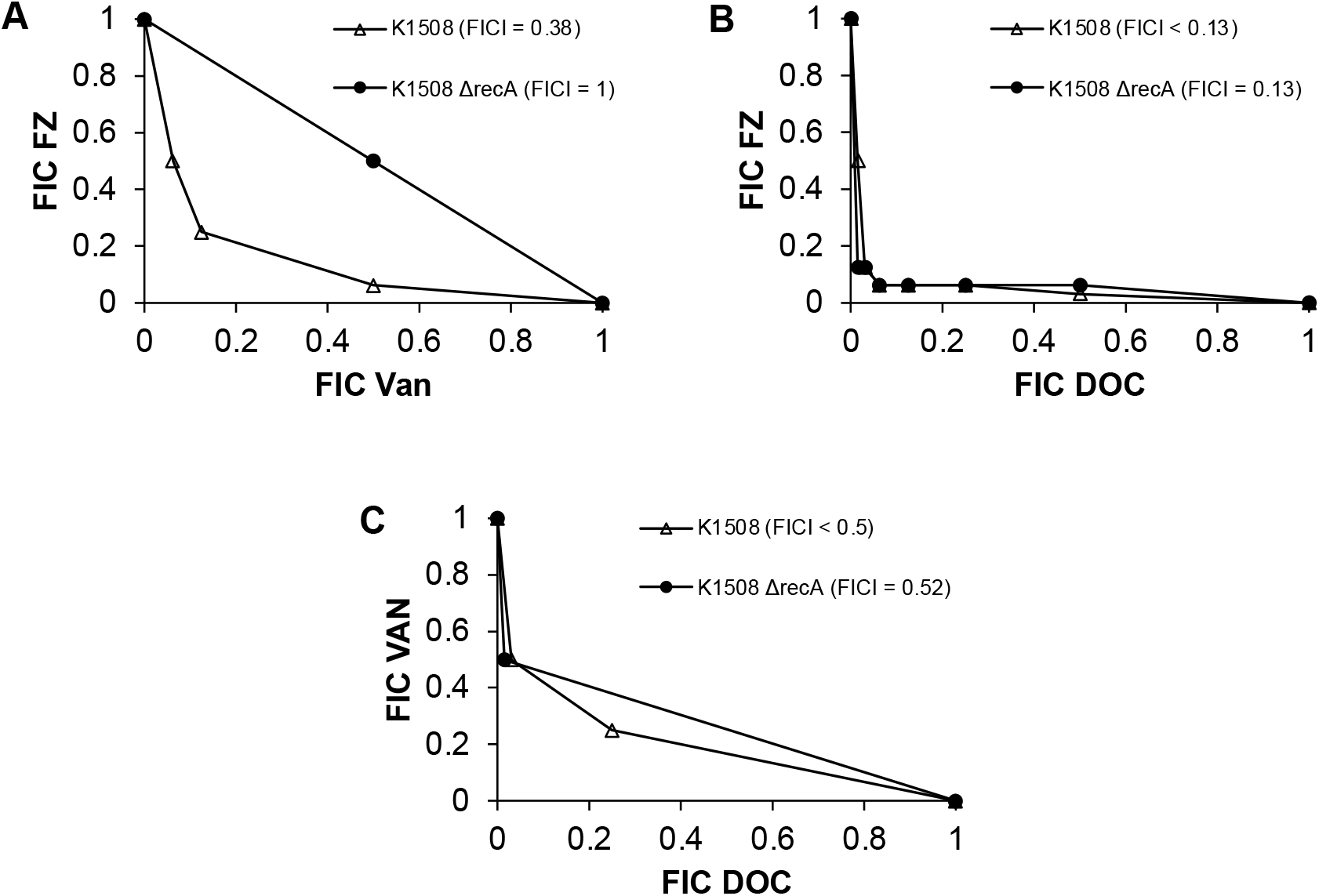
Effect of *recA* deletion on the two-drug interactions of (A) FZ and VAN, (B) FZ and DOC, and (C) VAN and DOC. Isobolograms were obtained using checkerboard assay and each data point represents fractional inhibitory concentrations (FIC; ratio of minimum inhibitory concentrations in combination *vs* alone).

## Discussion

Using a transcriptomics approach, the underlying mechanisms of action and synergy of FZ, VAN, and DOC, and the combination of all three (FVD), were investigated. The apparent similarity in the transcriptional responses induced by FZ and DOC in *E. coli* is likely the source of their synergy. This is in line with studies that found a higher likelihood of synergy occurring in drug combinations that induce either very similar or opposite gene perturbations (46, 47). In particular, FZ and DOC both resulted in the upregulation of genes encoding iron uptake systems and downregulation of those encoding iron storage and iron-utilizing proteins. This response is indicative of Fur inactivation, which usually occurs during iron starvation (29), and is consistent with previous reports of FZ and DOC increasing the expression of iron import genes (48, 49). An *E. coli fur* mutant grown in iron-rich conditions has been shown to result in a 2-fold iron decrease, while growth in the iron-depleted conditions showed a 14-fold reduction (50). Through measurements of the total intracellular iron levels, we determined a less than 2-fold intracellular iron decrease in the cultures containing FZ and DOC that corresponds to iron-rich conditions, thereby ruling out that these drugs cause real iron starvation.

The mechanisms by which FZ or DOC inactivate Fur and induce an iron starvation-like response are currently unknown. The mechanism may be as simple as reactive oxygen species oxidizing useable Fe^2+^ to unusable Fe^3+^ *via* the Fenton reaction (51) or damaging the Fe-S clusters (52). If so, iron import would be upregulated to supply iron to the labile iron pool and Fe-S cluster machinery. The Fur-Fe^2+^ complex has also been shown to be inactivated by nitric oxide (53). Incidentally, nitroheterocyclic drug reduction has been proposed to result in nitric oxide byproduct (54), though direct evidence for nitric oxide production during nitrofuran activation has not yet been reported. Despite the overall decrease of iron content in the cells, the inactivation of Fur will likely lead to an increase in the labile iron pool inside the cell that can increase oxidative damage and stress *via* the Fenton reaction (55).

FZ and DOC also caused gene perturbations usually observed in bacteriostatic translation inhibitors, such as downregulation of the central carbon metabolism and respiration, along with the upregulation of ribosomal proteins to compensate for the translational stress (44, 56, 57). Protein synthesis inhibition by nitrofurans has been reported and was proposed to be due to non-specific binding to ribosomal proteins and rRNA (13, 14). However, translation inhibition by DOC has not been demonstrated previously. Given that the Mg content of the cells is dramatically lowered by DOC, a possible connection between Mg homeostasis dysregulation by DOC and translational stress can be proposed: the decrease in total Mg levels by DOC will decrease the number of functioning ribosomes and thus inhibit translation in *E. coli* due to activation of the stringent response (58). However, due to the absence of (p)ppGpp regulation in the *E. coli* K1508 strain used in this work (*i.e*. *spoT* and *relA* mutant), translation control based on Mg^2+^ levels is possibly absent, which would lead to the unchecked upregulation of ribosome assembly even during Mg^2+^ deficiency, a phenomenon which has been demonstrated in *Salmonella* (58).

Our transcriptome analyses showed that FZ and DOC upregulated efflux pump genes (*acrA*, *acrB*), downregulated porin genes (*ompC*, *ompF*) (Fig. 2F), and induced stress responses that are expected to increase the tolerance to these agents. In this context, it seems contradictory that the combination is synergistic rather than antagonistic. A possible explanation for observed synergy is that some of these resistance mechanisms are somehow being inhibited by the combined action of FZ, VAN, and DOC. A proposed pathway for inhibition of these resistance mechanisms could be through Fur inactivation or downregulation of the central carbon metabolism. Both of these activities will have an inhibitory effect on the ETC (25, 59). FZ, DOC, and FVD downregulated the aerobic ETC genes *nuo* and *cyd*, which encode for two of the primary ETC complexes, NADH dehydrogenase I and cytochrome *bd*-I, that generate the proton motive force (PMF). These findings, along with the demonstrated overall decrease in aerobic respiration, could indicate diminished proton motive force in response to FZ and DOC. Since PMF is directly or indirectly required for the function of efflux pumps, low PMF is conducive for the accumulation of the antibacterials inside the cells by preventing efflux. These findings support our previous study in which deletion of *tolC*, or *acrA* efflux pump genes resulted in the loss or reduction of synergy between FZ and DOC, highlighting the importance of efflux in the synergy mechanism (7).

VAN treatment of *E. coli* did not induce much change in overall gene expression compared to the control. Even though only 17 genes were significantly differentially expressed, these genes do however give insight into the initial cellular effects of VAN. The majority of the 17 DEGs are members of envelope stress responses, particularly the Rcs pathway, which has been demonstrated to be induced by peptidoglycan-targeting antibiotics (40). Taken together, these findings indicate that VAN on its own can somehow cross the outer membrane of *E. coli*, although only in minimal amounts, and inhibit peptidoglycan synthesis. From the known mechanisms of action of VAN (peptidoglycan synthesis inhibitor) and nitrofurans (DNA damage), there does not seem to be an obvious connection as to why these drugs interact synergistically (8). One important mechanism of resistance to nitrofurans is the SOS response to DNA damage. Deletion of *recA*, making *E. coli* unable to mount an SOS response, resulted in the loss of synergy between FZ and VAN. Data presented here indicate SOS response’s involvement in the mechanism of synergy between FZ and VAN, possibly through inhibition of SOS induction by VAN, which can decrease the resistance to FZ’s DNA-damaging effect and therefore increase lethality. The *recA* deletion also increased the FICI for the triple combination, but the interaction is still synergistic, supporting the concept of multiple interaction points that contribute to the synergy of FVD.

This study searched for the possible mechanistic bases of the synergy between FZ, DOC, and VAN using transcriptomics and biochemical approaches. The transcriptional responses shed light on the modes of action of FZ and DOC. By analysing their highly similar perturbed pathways, in combination with VAN effects, it was possible to propose mechanisms for the synergy based on identified changes in gene expression. Particularly, we propose that the combined effects on the Fur pathway lead to (1) an increased labile iron pool that can cause oxidative damage to proteins and DNA and (2) inhibition of ETC that can diminish PMF and subsequently inhibit PMF-dependent efflux activity. FZ and DOC’s combined translational stress and VAN’s possible inhibition of the SOS response, which can amplify the DNA damage, are also possible sources of synergy. FZ, DOC, and VAN affected correlated pathways that likely result in the suppression of resistance mechanisms and amplification of damaging effects. Although further work is warranted to fully elucidate these mechanisms, this study lays the groundwork for the development of this combination into a viable clinical therapy for tackling multidrug-resistant bacterial infections.

## Materials and Methods

### Bacterial strain, growth conditions, and checkerboard assay

*E. coli* K1508 (MC4100 [*F*^−^ *araD*^−^ *Δlac* U169 *relA1 spoT1 thiA rpsL* (Str^R^)] *ΔlamB106*) (20) was grown aerobically in 2xYT medium at 37 °C. *kan^R^ recA* deletion mutation from the Keio collection (60) were introduced into *E. coli* K1508 using phage P1 transduction, as previously described (61). To eliminate potential polar effects on downstream genes in the operon, the FLP recombinase recognition target (FRT)-flanked *kan^R^* cassette was excised using FLP-mediated recombination using plasmid pCP20 (62).

### Determination of MIC and checkerboard assay

MIC determinations and checkerboard assays were performed using the broth microdilution method in a 384-well plate according to the CLSI guidelines (63), with minor changes. 2xYT medium was used, the inoculum concentration is 1 × 10^6^ CFU/mL, the plates were incubated at 37 °C for 24 h, and all experiments were performed in triplicate. MIC is the lowest concentration that completely inhibits growth.

Synergistic, antagonistic, and no interactions were determined using the fractional inhibitory concentration index (FICI) method, using the equation:

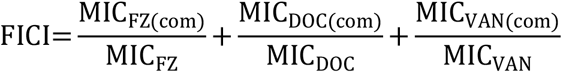

where MIC_FZ(com)_, MIC_DOC(com)_, and MIC_VAN(com)_ is the MIC of FZ, DOC or VAN when used in combination and MIC_FZ_, MIC_DOC_, and MIC_VAN_ is the MIC when used alone. Using the lowest FICI, the interactions were interpreted as synergistic if FICI ≤ 0.5; indifferent if 0.5 < FICI ≤ 4.0 and antagonistic if FICI > 4.0 (64).

### Total RNA isolation and sequencing

The concentrations that gave 50% growth inhibition (IC_50_) at 24 h was chosen for the transcriptomics study. The IC_50_ determination was performed in 384-well plates on *E. coli* K1508 at 1 × 10^6^ CFU/mL in a total volume of 50 μL. The plate was incubated at 37 °C for 24 h, and optical density at 600 nm (OD600) was determined. The mean % growth inhibition was calculated and the R package drc v3.0-1 (65) was used to plot the concentration-response (% inhibition) curves fitted with a four-parameter log-logistic model to determine the IC_50_. For the combination, the most synergistic combination (*i.e*. lowest FICI) in a checkerboard assay were first determined (Fig. S2), then fixed-ratio dilutions of these concentrations were used to plot a concentration-response curve. The IC_50_ for each antibacterial and combination, which was used in all the assays, are summarized in Table S1.

Exponentially growing cultures of the *E. coli* K1508 at 5 × 10^7^ CFU/mL were treated with IC_50_ antibacterial(s) in a final volume of 25 mL. The DMSO concentration for all treatments was fixed at 0.1%. After incubation at 37 °C with shaking at 200 rpm for 1 h, the cultures were harvested by centrifugation. The resulting pellet was then resuspended in 1 mL of resuspension buffer (20 mM sodium acetate pH 5.5, 1 mM EDTA, 1% SDS) and homogenized by beat beating. The samples were then subjected to phenol-chloroform nucleic acid extraction, as previously described (66), except acid phenol with pH of 4.45 was used to extract RNA.

Experiments were conducted in quadruplicate, and the samples were sent to Novogene Co., Ltd. (Beijing, China) for rRNA depletion using a Ribo-Zero^™^ rRNA Removal Kit (Illumina), library preparation using the NEBNext^®^ Ultra^™^ Directional Library Prep Kit for Illumina^®^ (New England Biolabs, USA), and subsequent 150-bp paired-end RNA sequencing on a HiSeq 2500 sequencer (Illumina).

### RNA-Seq analysis

Before analysing the data, the quality of the reads was checked using FastQC v0.11.7-5 (67). The RNA sequencing reads were then mapped against the *E. coli* K1508 genome (NCBI GenBank accession CP072054) using HISAT2 v2.1.0 (68) and the number of reads that mapped to a gene was counted using featureCounts v1.6.0 (69) (Table S2).

Differential expression analysis was carried out using DESeq2 v1.26.0 (70). To better represent effect size (gene expression), log_2_FC estimates were shrunk using the apeglm v1.8.0 shrinkage estimator (71). Differentially expressed genes (DEG) were defined as genes with an adjusted p-value (multiple test adjustment using the Benjamini-Hochberg method) of less than 0.01 (adj p < 0.01) and fold changes greater than 1.5 (|log_2_FC| > 0.58). The DEGs were clustered using k-means clustering of regularised-log transformed normalised counts into optimal k number of clusters identified by Mclust function of the R package, Mclust v5.4.6 (72). Gene Ontology (GO) term enrichment of the DEGs was performed using statistical overrepresentation test (Fisher’s exact test with Benjamini-Hochberg false discovery rate (FDR) multiple test correction) in PANTHER v15.0 (73). The significantly overrepresented GO terms were selected using an FDR cutoff of 0.05.

### Mutagenicity Assay

Mutation frequencies were measured as described previously (74). Briefly, exponentially-growing *E. coli* K1508 at 1 ×10^7^ CFU/mL were treated with IC_50_ FZ, VAN, and DOC, alone and in combination, in a final volume of 10 mL in 2xYT medium. The cultures were incubated at 37 °C with shaking at 200 rpm for 24 h. The cultures were then centrifuged at 5000 ×g for 10 min and resuspended in maximum recovery diluent (0.1% peptone, 0.85% NaCl). Serial dilutions were plated in triplicate onto 2xYT agar containing 100 μg/mL rifampicin to select for rifampicin-resistant colonies and on non-selective 2xYT agar to count the total number of colonies. The plates were scored after 24 h at 37 °C. The mutation frequency was calculated by dividing the number of rifampicin-positive colonies by the total number of colonies from 9-11 biological replicates.

### Oxygen consumption

Oxygen consumption was measured as previously described (75). Briefly, *E. coli* K1508 culture at OD600 of 0.3 was treated with IC_50_ FZ, VAN, and DOC, alone and in combination, at 37 °C for 1 h. Cells were then diluted in air-saturated 2xYT to OD600 of 0.2, and dissolved oxygen was measured in a closed chamber with constant stirring using a Clark-type oxygen electrode (Rank Brothers Ltd.) linked to a chart recorder (Vernier LabQuest Mini).

### Metal concentration by ICP-MS

Antibiotic-treated *E. coli* cultures in a total volume of 80 mL were processed the same way as for the transcriptomics analyses. After antibiotic treatment, cells were collected and prepared for ICP-MS, as previously described (76). Briefly, cells were harvested by centrifugation (5000 ×g, 10 min), then washed twice with 25 mL phosphate-buffered saline (PBS) containing 0.5 mM EDTA, then twice with PBS. All samples were adjusted to a cell number of 2 × 10^9^ CFU based on their OD600 values. Washed cell pellets were then digested with 500 μL of 70% (wt/vol) nitric acid (≥99.999% trace metals basis) at 80 °C overnight. Each sample was diluted 1:20 in Milli-Q water (18.2 MΩ), giving a final acid matrix of 3.5%. The samples were then sent to the University of Waikato Mass Spectrometry Facility to analyze metal content by ICP-MS on an Agilent 8900 system.

## Data availability

The transcriptomic raw data were deposited in GenBank under BioProject accession PRJNA642878.

## Acknowledgements

We are grateful to Dr. Van Hung Vuong Le for sequencing the genome of *E. coli* K1508, to Adrian Koolaard for technical help with the oxygen electrode, and to Dr. Amanda French of University of Waikato for carrying out the ICP-MS analysis.

## Supplemental Material Legends

**Fig. S1** Overall transcriptomic changes of *E. coli* K1508 in response to FZ, VAN, DOC, or the FVD combination. (A) Principal component analysis plot showing the correlation and variation between and within treatment groups. (B) Volcano plots showing the significantly upregulated and downregulated genes (red points, adj p < 0.01 and |log_2_FC| > 0.58) in each treatment group. The ten most upregulated, downregulated, and statistically significant genes are labeled. (C) Venn diagram of differentially expressed genes.

**Fig. S2** GO term enrichment analysis of the significantly differentially expressed genes in VAN-treated *E. coli* from the O’Rourke et al. dataset (44). Biological process GO terms overrepresented in the significantly (A) upregulated (B) and downregulated genes. GO terms associated with cell envelope were highlighted in blue, molecule transport in green, and SOS response in orange.

**Fig. S3** Interaction of FZ, VAN, and DOC in the growth inhibition of (A) *E. coli* K1508 and (B) K1508 Δ*recA* mutant. Each data point represents minimal inhibitory concentrations (MIC) identified by checkerboard assay. The most synergistic concentration combinations, which gave the lowest FICI are highlighted as follows: red for FZ+VAN+DOC, blue for FZ+DOC, green for FZ+VAN, and orange for VAN+DOC. The MIC of DOC cannot be determined for *E. coli* K1508 since solubility is limited to 80 mg/mL, which was used as proxy to calculate FICI, and therefore actual FICI will be smaller than the calculated FICI.

**Table S1** Concentrations of FZ, VAN, and DOC used in the study

**Table S2** Quality of reads and read alignment statistics

**Dataset S1** Differential expression analysis of *E. coli* K1508 genes after treatment with FZ, VAN, DOC, or the FVD combination

**Dataset S2** Biological process GO term enrichment analysis of differentially upregulated and downregulated genes in each of the treatments

**Dataset S3** k-means clustering and subsequent GO term enrichment analysis of all the differentially expressed genes across treatments

